# Gq-pathway activation in hippocampal CA1 astrocytes rescues ischemia-induced memory deficits and synaptic plasticity

**DOI:** 10.64898/2026.03.20.713091

**Authors:** Yujie Chen, Lei Wang, Jiayao Wang, Yutong Zhou, Hui Shen, Yuguang Wang, Wei Qin, Bo Liu, Baihe Chen, Yiting Huang, Wenting Guo, Huisen Xu, Qian Tian, Chen Zheng

## Abstract

More than 30% of stroke survivors develop post-stroke cognitive impairment (PSCI), for which current neuron-centric therapies not only lack cellular specificity but also carry risks of adverse effects such as epilepsy. Here, we explore the therapeutic potential of astrocytes by chemogenetically activating Gq signaling in hippocampal astrocytes, which rescues memory deficits and synaptic plasticity impairments in a mouse model of ischemic stroke. Gq activation restores dendritic complexity, spine density, and long-term potentiation in hippocampal CA1 neurons. Fiber photometry further reveals that astrocytic Ca²⁺ signals precede neuronal activity by 600 ms during novel environment exploration, indicating that astrocytes prime memory encoding. In contrast, Gi pathway activation induces pathological neuronal hyperactivity without cognitive improvement. These findings establish that astrocytes regulate post-stroke recovery through pathway-specific calcium signaling and uncover a previously unknown temporal hierarchy in astrocyte-neuron communication during memory processing, offering a new glia-targeted strategy to overcome the limitations of current neuromodulation approaches for PSCI.

## Introduction

Post-stroke cognitive impairment (PSCI) is a leading cause of disability in stroke survivors, yet effective treatments remain critically lacking. Currently available neuron-targeted interventions, such as trans-cranial magnetic stimulation and deep brain stimulation, have shown short-term cognitive improvement in PSCI^1, 2^. However, their long-term efficacy is often limited, and more critically, they carry the risk of inducing seizures^3, 4^. Given that conventional therapeutic strategies primarily focus on modulating neuronal activity and typically lack cellular specificity, their clinical application is further constrained by insufficient sustained benefits and potential adverse effects. In this context, astrocytes—the most abundant glial cells in the central nervous system—have emerged as a promising therapeutic target due to their essential role in synaptic plasticity, memory formation and post-stroke rehabilitation^3, 5^.

Astrocytes actively participate in neural information processing through the “ tripartite synapse”: they sense neuronal activity, respond via calcium signaling, and release gliotransmitters to precisely modulate synaptic function^6^. Adamsky et al. demonstrated that physiological astrocytic Ca²⁺ signaling promotes memory allocation in vivo^3^. Importantly, recent research demonstrates that activated astrocytes form functional ensembles during learning — termed learning-associated astrocytes (LAAs) — which can be reactivated during memory retrieval to enhance recall efficiency^7^. Notably, direct neuronal activation may impair contextual memory, suggesting that astrocyte-mediated modulation offers superior physiological precision. Following ischemic stroke, astrocytic microdomain Ca²⁺ transients in the peri-infarct region are significantly reduced^8^. Conversely, optogenetic precision modulation of astrocytic calcium signaling has been shown to enhance motor recovery after chronic capsular infarct^5^. Moreover, astrocytes exhibit remarkable ischemic tolerance and can transfer functional mitochondria to compromised neurons via calcium-dependent mechanisms, providing neuroprotection^9^.These findings collectively indicate that modulating astrocytic calcium dynamics represents a promising strategy for promoting post-stroke recovery.

Nevertheless, critical questions remain: Can astrocytic calcium signaling regulate the pathogenesis and functional recovery of PSCI? What are the underlying mechanisms? To address these questions, we combined chemogenetic activation with synaptic analyses, behavioral testing, and in vivo fiber photometry to systematically investigate the role of hippocampal astrocytic Ca²⁺ signaling in PSCI. We demonstrate that Gq pathway activation in CA1 astrocytes rescues memory deficits in a mouse model of ischemic stroke and restores synaptic plasticity indicators such as dendritic complexity, spine density, and long-term potentiation. Furthermore, we reveal a precise temporal coordination between astrocytic and neuronal activity during memory processing, providing novel theoretical insights and a strategic framework for astrocyte-targeted therapy in PSCI.

## Results

### Cerebral ischemia impairs spatial reference memory

To evaluate the impact of cerebral ischemia on spatial memory, we performed the Y-maze novelty preference test (NPT) in mice 3 days after inducing transient global cerebral ischemia through bilateral common carotid artery ligation (BCAL) for 1 hour (Fig. 1A). In this behavioral paradigm, mice were first allowed to explore two arms of the maze for 15 minutes, followed by a 1-hour interval before being reintroduced to the maze with all three arms accessible. Both BCAL and sham-operated mice showed comparable levels of general locomotor activity and exploration, as evidenced by similar total exploration distances and time spent in the start arm (Fig. 1B). All animals demonstrated a natural preference for the novel arm over the familiar arm (Fig. 1C). However, ischemic mice exhibited significantly weaker novelty preference compared to sham controls: novel arm exploration distance was reduced from 66.76 ± 8.16% in sham mice to 57.05 ± 5.17% in BCAL mice (P < 0.01), while novel arm exploration time decreased from 68.97 ± 10.85% to 56.81 ± 8.97% (P < 0.01) (Fig. 1D and E). These findings suggest certain deficits in spatial reference memory in mice after ischemic stroke.

**Fig. 1.**
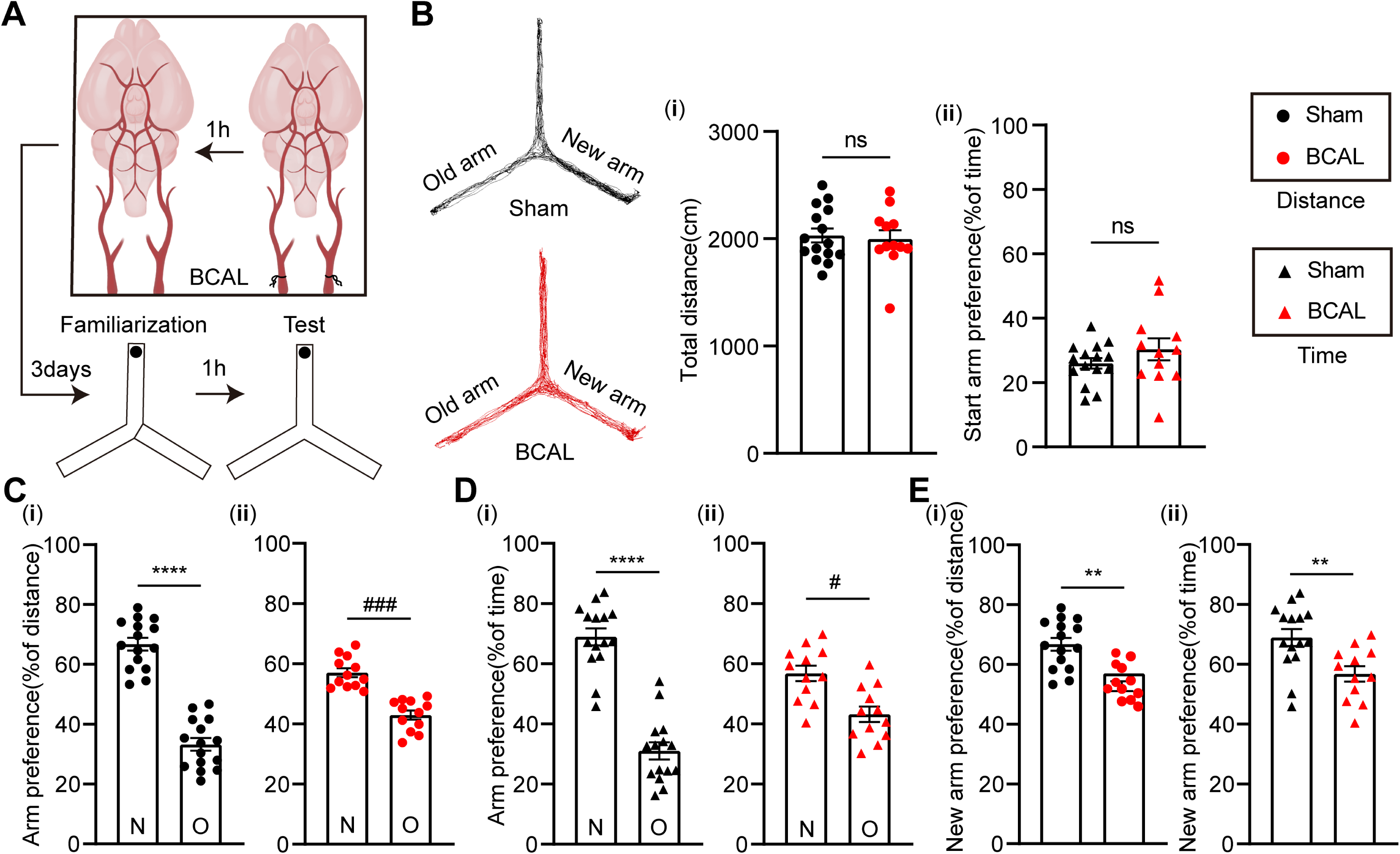
Cerebral ischemia impairs novelty preference in Y-maze test. (A) Experimental timeline of Y-maze novel place test (NPT) following cerebral ischemia. (B) Left: Representative movement traces. Right: total distance (i) and start arm exploration time (ii) (sham vs BCAL). (C) Novel vs familiar arm exploration distance in sham (i) and BCAL (ii) mice. (D) Novel vs familiar arm exploration time in sham (i) and BCAL (ii) groups. (E) Novel arm preference (distance (i) and time (ii)) comparison between groups (sham: n = 15 mice; BCAL: n = 12 mice). N, new arm; O, old arm. Data: mean ± SEM. Statistics: \*\**P* < 0.01, \*\*\*\**P* < 0.0001 vs sham; ^#^*P* < 0.05, ^###^*P* < 0.001 vs BCAL; ns, not significant. Independent t-test (B, E); paired t-test (C, D).

### Astrocytic activation rescues spatial reference memory deficits

Based on established roles of hippocampal astrocytes in memory, we tested whether their activation during Y-maze exploration could rescue memory deficits. Mice were divided into three groups receiving CA1 injections of: hM3D (Gq), hM4D (Gi), or mCherry (control) viruses. Following 3 weeks of viral expression and BCAL surgery, memory was assessed via NPT (Fig. 2A and B).

**Fig. 2.**
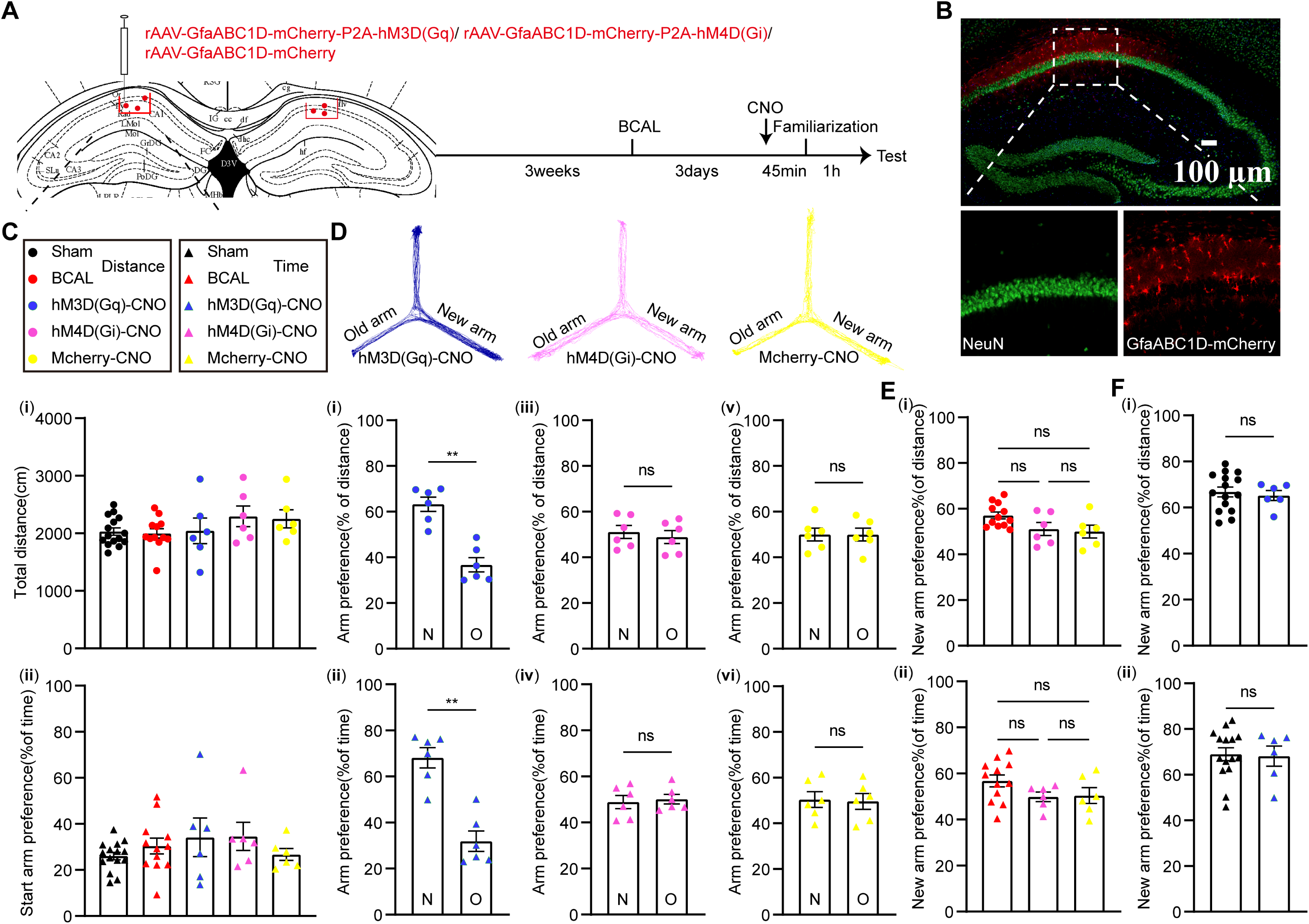
Gq-mediated astrocytic activation rescues novelty preference deficits after global cerebral ischemia. (A) Experimental timeline showing viral injection in CA1 and behavioral testing. (B) Schematic of CA1-specific mCherry expression via GfaABC1D promoter. (C) Total distance (i) and time spent in start arm (ii) across groups (sham, BCAL, hM3D, hM4D, mCherry). (D) Top: representative Y-maze traces. Bottom: Novel (N) vs familiar (O) arm exploration in hM3D (i, ii)-, hM4D (iii, iv)-, and mCherry (v, vi)-expressing mice. (E) Novel arm preference (distance (i) /time (ii)) in hM4D vs mCherry vs BCAL. (F) Novel arm preference (distance (i) /time (ii)) comparison between sham and hM3D groups (n = 6 mice/group). Data: mean ± SEM. \*\**P* < 0.01 vs hM3D; ns = not significant. Statistics: one-way ANOVA (C, E); paired t-test (D); independent t-test (F).

To test the effect of astrocytic activity on memory performance, CNO was administered 45 min before Y-maze test. No significant effect of CNO administration on total exploration distance was observed (Fig. 2C). Astrocytic activation of hM3D (Gq) with CNO resulted in a significant elevation in the exploration distance of the novel arm compared to the familiar arm (63.25 ± 7.724%), but the hM4D(Gi) group (51.13 ± 6.917%) and the control group (50.01 ± 6.111%) didn’t show a novelty preference, because they explored two arms of a maze at a similar level (Fig. 2D). Compared with the BCAL group, Gi-coupled hM4D with CNO application did not improve any arm preference (Fig. 2E). However, astrocytic chemogenetic activation of Gq pathway rescued spatial preference in BCAL mice, reaching levels similar to sham controls (Fig. 2F). Similarly, the CNO application had no effect on explored willingness (Fig. 2C). Furthermore, activation of hM3D (Gq) increased the percentage of exploration distance as well as increased the percentage of exploration time (68.12 ± 10.77%) during NPT (Fig. 2D). These findings indicate that Gq-mediated astrocytic activation confers its cognitive-enhancing effects to improve memory impairment after ischemic stroke and supports the potential therapeutic effect.

To confirm that the observed cognitive benefits were specifically attributable to chemogenetic activation of astrocytic signaling rather than non-specific effects of viral injection, we included saline-injected control groups. As shown in Supplemental Figure 1B–C, saline administration failed to rescue novel arm preference deficits in mice expressing either Gq- or Gi-coupled chemogenetic receptors, confirming that the behavioral effects were specifically driven by chemogenetic activation.

### Effect of astrocytes on synaptic plasticity

#### Gq-mediated astrocytic activation drives post-ischemic synaptic remodeling

To investigate synaptic structural plasticity underlying memory impairment, we performed Golgi-Cox staining and quantitative sholl analysis in hippocampal CA1 neurons. Compared to sham controls (158.7 ± 40.90%), ischemic injury (BCAL) significantly reduced dendritic complexity (107.1 ± 29.65%, P < 0.05) (Fig. 3A and 3C). Remarkably, chemogenetic activation of astrocytes via hM3D(Gq) not only restored dendritic complexity to 177.1 ± 56.34% (Fig. 3C) but also normalized spine density (10.29 ± 2.162% vs BCAL 7.732 ± 1.608%), reaching levels comparable to sham controls (10.82 ± 2.150%) (Fig. 3D). In contrast, chemogenetic manipulation via hM4D(Gi) failed to ameliorate these structural deficits. This functional dissociation suggests that astrocytic Ca²⁺ signaling operates through pathway-specific mechanisms, where Gq activation is both necessary and sufficient for synaptic remodeling after ischemic injury.

**Fig. 3.**
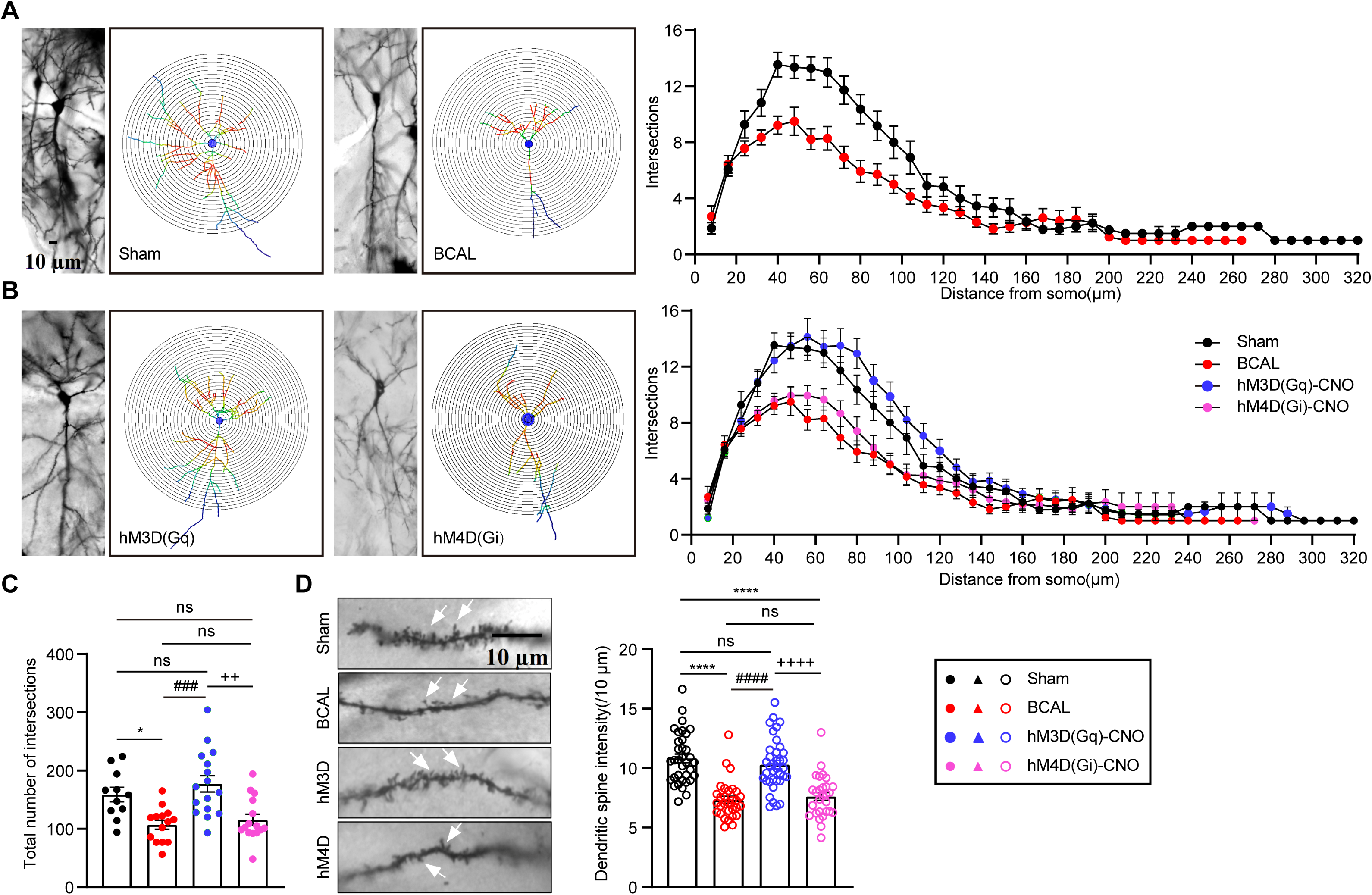
Pathway-specific effects of astrocytic modulation on CA1 neuronal structure after ischemia. (A) Golgi-stained CA1 neurons and sholl analysis in sham vs BCAL groups. (B) Neuronal complexity in hM3D- vs hM4D-expressing mice post-CNO. (C) Intersection number comparison across groups. (D) Left: representative dendritic spines. Right: spine density quantification (sham:11neurons/9slices from 1 mice; BCAL:14neurons/7slices from 1mice; hM3D:16 neurons/13slices from 1 mice; hM4D:15 neurons/11slices from 1 mice). Data: mean ± SEM. \**P* < 0.05, \*\*\*\**P* < 0.01 vs sham; ^###^*P* < 0.001, ^####^*P* < 0.0001 vs BCAL; ^++^*P* < 0.01, ^++++^*P* < 0.0001 vs hM3D. Statistics: one-way ANOVA (C, D).

#### Gq-mediated astrocytic activation enhances synaptic transmission in the Schaffer collateral pathway

Given the critical dependence of neuronal function on morphological integrity, which is compromised by ischemia and modulated by astrocytes, we investigated astrocyte-mediated regulation of synaptic transmission in hippocampal CA1. Using acute slice electrophysiology, we assessed long-term potentiation (LTP) at Schaffer collateral pathway through local field potential recordings under artificial cerebrospinal fluid (ACSF) conditions (Fig. 4A). High-frequency stimulation (HFS) by 4 stimulus trains of 100 pulses (1s duration, 10s intervals, 100Hz) induced robust LTP in sham controls at 60 min post-HFS (155.5 ± 23.60% baseline). This plasticity was significantly impaired following cerebral ischemia (108 ± 20.34% baseline; Fig. 4B and 4C). Notably, chemogenetic activation of astrocytes via hM3D(Gq) completely rescued LTP impairment (144.3 ± 21.7% of baseline; Fig. 4D and 4E). In contrast, hM4D(Gi) manipulation showed no significant effect on ischemic LTP (100.2 ± 13.83% of baseline; Fig. 4D-F), suggesting that Gi-coupled pathways may not sufficiently modulate the astrocytic mechanisms supporting synaptic potentiation in this paradigm.

**Fig. 4.**
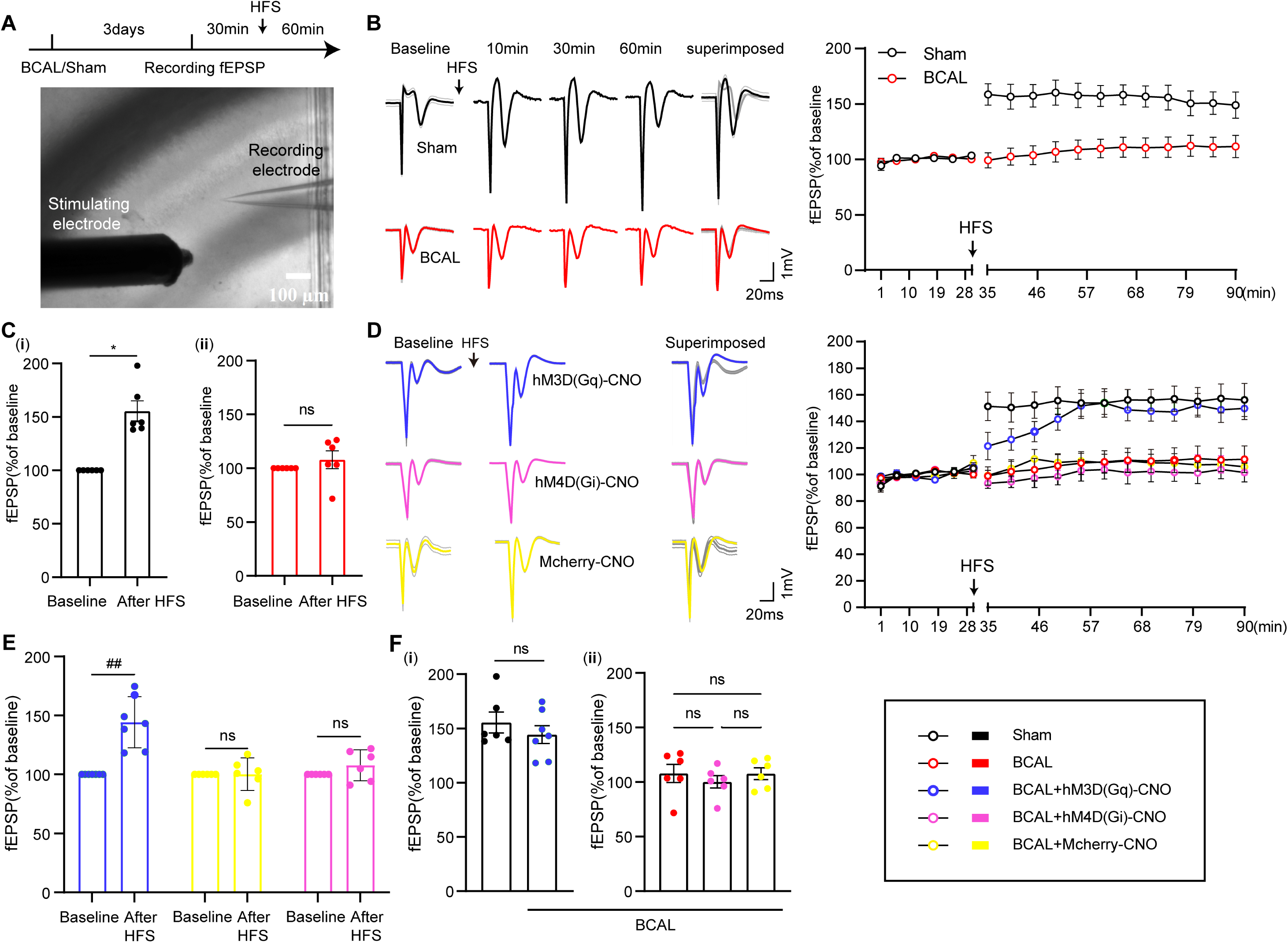
Gq-mediated astrocytic activation rescues ischemia-induced LTP deficits in hippocampal CA1. (A) Experimental timeline and LTP induction protocol in Schaffer collaterals. (B) Left: representative fEPSP traces pre- and post-HFS (10/30/60 min). Right: LTP quantification. (C) Mean fEPSP (last 30 min) in sham (i) and BCAL (ii) groups. (D) Left: fEPSP traces pre-/post-HFS. Right: LTP comparison across groups. (E) Mean fEPSP in chemogenetic groups. (F) fEPSP changes (hM3D vs sham (i); hM4D/mCherry vs BCAL (ii)). hM3D: n = 7 slices from 7 mice; other groups: n=6 slices from 6 mice. Data: mean ± SEM. *P < 0.05 vs sham; ^##^*P* < 0.01 vs hM3D. Statistics: paired t-test (C, E); unpaired t-test (F, i); one-way ANOVA (F, ii).

#### Divergent modulation of post-ischemic neuronal activity by astrocytic Gq versus Gi signaling pathways

To assess region-specific neural activity patterns following ischemia and astrocytic modulation, we quantified c-Fos expression as a marker of neuronal activation. In the Y-maze spatial memory test (Fig. 5A), ischemic (BCAL) mice showed significantly reduced neuronal c-Fos⁺ nuclei in CA1 compared to sham controls (BCAL: 2.773 ± 1.326% vs Sham: 8.552 ± 2.955% cells/mm², P = 0.0118), while astrocytic c-Fos levels remained unchanged (BCAL: 0.3242 ± 0.3525% vs Sham: 0.3158 ± 0.3679% GFAP+ cells/mm², P = 0.9746) (Fig. 5B and 5C).

**Fig. 5.**
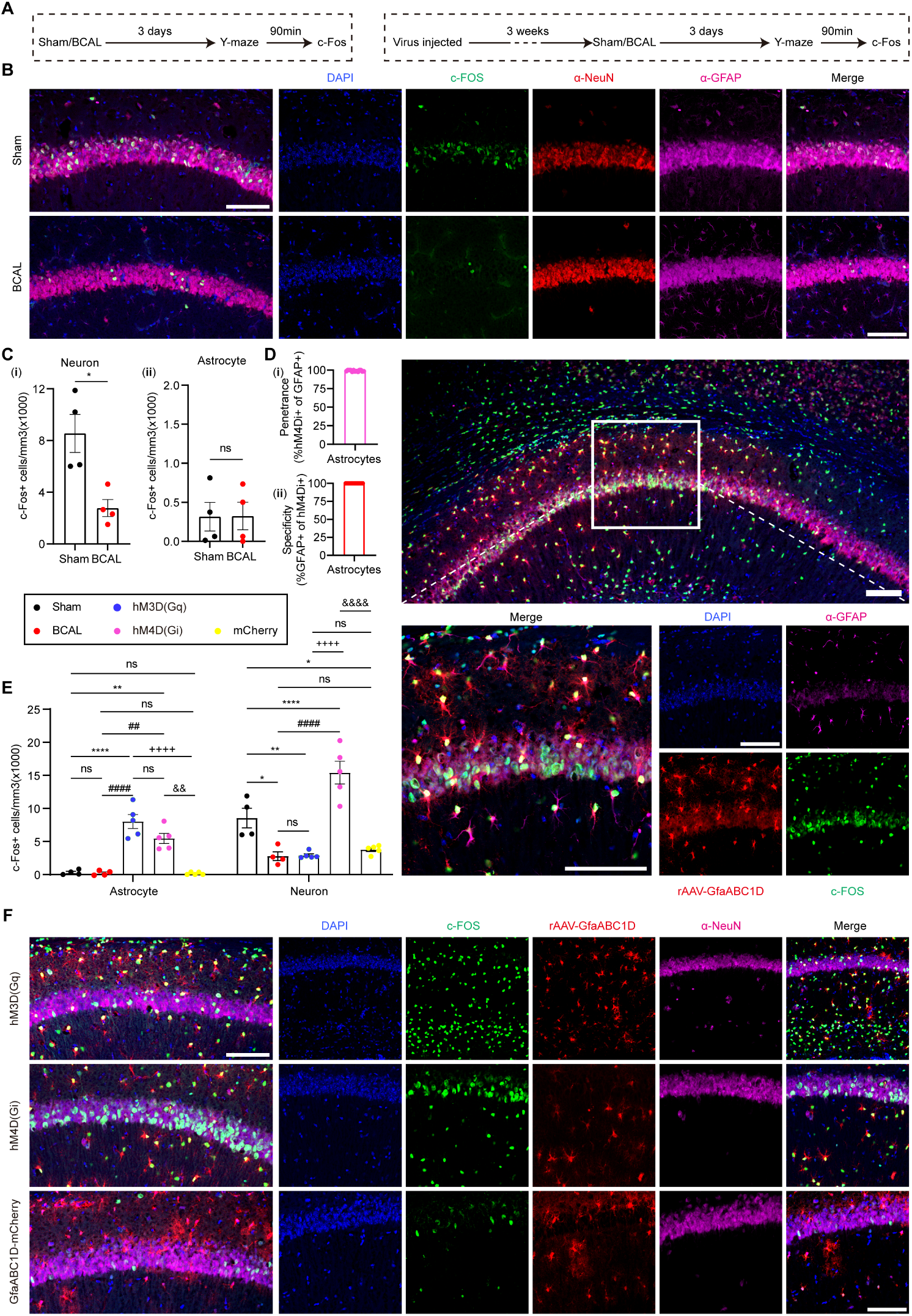
Gq-mediated astrocytic activation modulates memory-associated neuronal recruitment. (A) Experimental timeline. (B) Representative CA1 images showing c-Fos^+^/NeuN^+^ neurons and GFAP+ astrocytes in sham vs BCAL groups. (C) Quantified c-Fos intensity in NeuN^+^ (i) and GFAP^+^ (ii) cells. (D) Left: hM4D expression specificity (>97% astrocyte targeting, 884/894 cells from 3 mice). Right: confocal images of hM4Di/GFAP/c-Fos colocalization in chemogenetic groups. (E-F) c-Fos intensity in NeuN^+^/GFAP^+^ cells across groups. Sample sizes: sham/BCAL (17-20 slices from 4 mice); chemogenetic groups (16-24 slices from 5 mice). Data: mean ± SEM. \**P* < 0.05, \*\**P* < 0.01, \*\*\*\**P* < 0.0001 vs sham; ^##^*P* < 0.01, ^####^*P* < 0.0001 vs BCAL; ^++++^*P* < 0.0001 vs hM3D; ^&&&&^P < 0.0001 vs hM4D. Statistics: unpaired t-test (C); two-way ANOVA (E). Scale bar: 100 μm.

Using astrocyte-specific chemogenetic tools, we found mCherry expression was restricted to astrocytic membranes with high transduction efficiency (> 97% GFAP⁺ cells expressed mCherry) and specificity (> 99% hM4Di⁺ cells were GFAP⁺) (Fig. 5D and Fig. S1A). Notably, Gq pathway activation (hM3D + CNO) significantly increased astrocytic c-Fos (Gq: 8.029 ± 2.352% cells/mm² vs mCherry: 0.1490 ± 0.2044% cells/mm², P < 0.0001) without altering neuronal c-Fos levels (Gq: 2.903 ± 0.5253% cells/mm² vs BCAL: 2.773 ± 1.326% cells/mm², P > 0.9999). Strikingly, while Gi pathway manipulation (hM4D + CNO) produced comparable astrocytic activation (5.467 ± 1.700% cells/mm²), it triggered disproportionate neuronal hyperactivation (15.42 ± 3.853% vs BCAL, P < 0.0001), that exceeded even sham control levels (Fig. 5E and 5F). This dissociation between astrocytic and neuronal responses suggests Gi-coupled receptors may engage distinct signaling cascades in the ischemic microenvironment. The pathological nature of this hyperactivation was further evidenced by its coexistence with persistent cognitive impairment, indicating a breakdown in normal hippocampal encoding mechanisms where elevated c-Fos failed to translate to functional recovery.

#### Astrocyte–neuron communication enables precision control of hippocampal Ca²⁺ oscillations in post-stroke cognitive recovery

To further investigate the real-time astrocytic and neuronal Ca^2+^ dynamics during recall in NPT, the genetically encoded Ca^2+^ indicator (GECI)-based fiber photometry was coupled with the NPT to monitor astrocytic and neuronal activity in the CA1 of mice, respectively. In this task, all mice were divided into the astrocyte group, which was injected with a green astrocytic Ca^2+^ indicator: rAAV-GfaABC1D-cyto-GCaMP6f-SV40 (GfaABC1D-GCaMP6f) and the neuron group, which was injected with a green neuronal Ca^2+^ indicator: rAAV-hSyn-GCaMp6f-WPRE-hGH (hSyn-GCaMp6f) (Fig. 6A). Within the virally transduced region, astrocytic GCaMp6f expression was limited to the astrocytic cytoplasm (> 97% GCaMp6f positive cells were also GFAP positive) (Fig. S2C and S2D). In contrast, neuronal GCaMp6f expression was limited to neuronal membranes, similar to that of the neuronal marker NeuN (> 97% colocalization) (Fig. S2B and S2D). We observed that both astrocytic and neuronal Ca^2+^ exhibited specific patterns responding to the NPT.

**Fig. 6.**
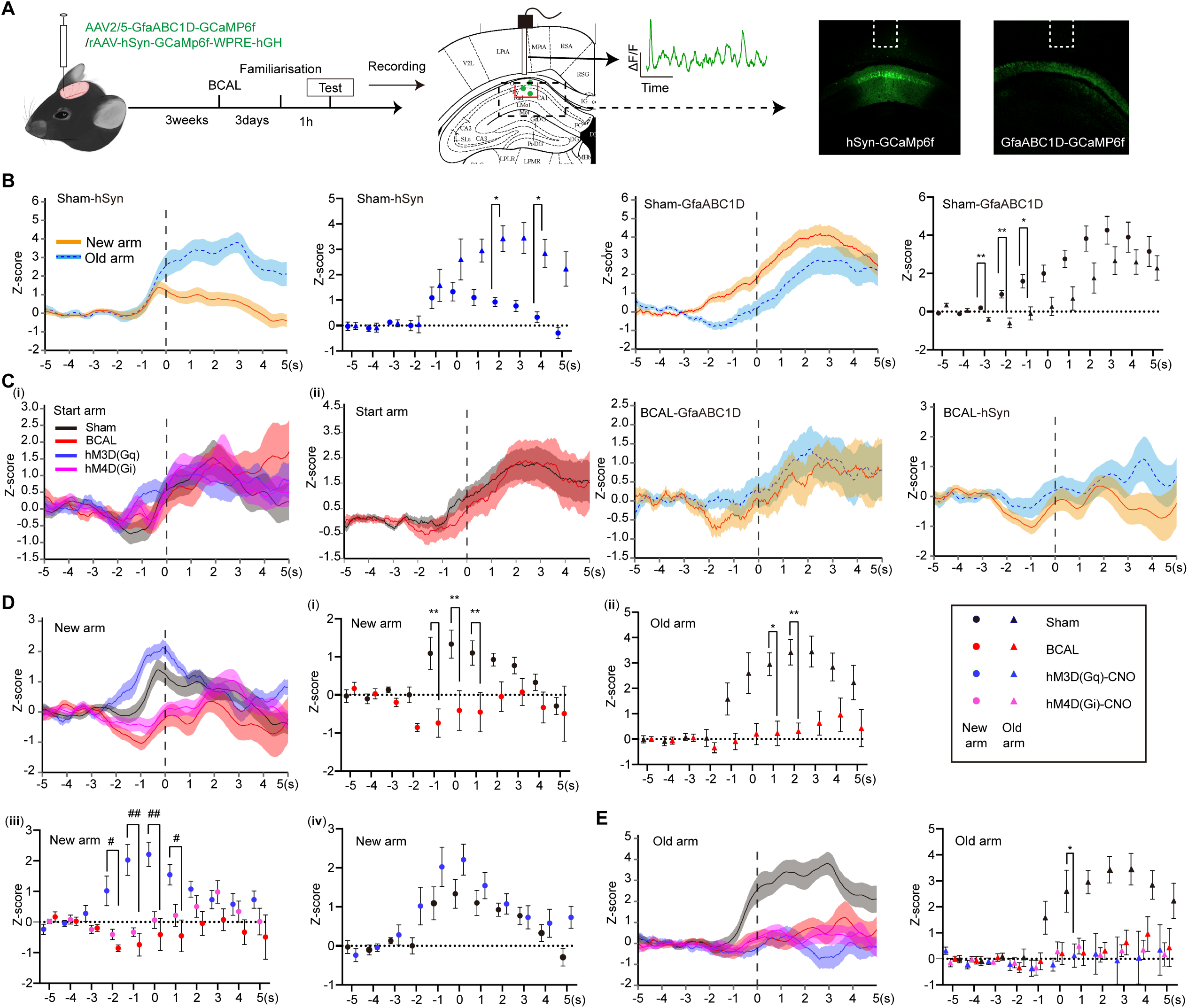
Astrocytic Ca²⁺ dynamics guide neuronal activity during spatial decisions. (A) Left: experimental timeline and fiber photometry setup. Right: CA1 viral expression (white box: fiber location). (B) Arm-selective Ca^2+^ responses in sham mice (neurons: hSyn; astrocytes: GfaABC1D). (dashed line: maze entry). (C) Left: Baseline Ca^2+^signals in neurons (i) and astrocytes (ii) during start-arm approaches. Right: BCAL-induced disruption of arm-selective responses. (D-E) Quantitative comparison of neuronal Ca^2+^ changes (sham vs BCAL) (i, ii). Chemogenetic modulation effects during novel (iv) and familiar (E) arm choices. Sample sizes: sham (n = 15 mice: 9 for GfaABC1D-GCaMP6f, 6 for hSyn-GCaMP6f); BCAL (n = 12 mice: 6/group); chemogenetic (n = 6 mice/group). Data: mean ± SEM. \**P* < 0.05, \*\**P* < 0.01 vs sham; ^#^*P* < 0.05, ^##^*P* < 0.01 vs hM3D. Statistics: two-way ANOVA.

Fiber photometry recordings with cell-type-specific GCaMP6f revealed that in sham-operated mice, both astrocytic and neuronal Ca²^+^ signals significantly increased as mice approached the center of the Y-maze (astrocytes: Z-score = 0.001328 ± 0.8876% to peak 2.328 ± 2.626%; neurons: -0.5917 ± 0.6997% to peak 1.360 ± 1.218%; both P < 0.01 vs baseline). Notably, neuronal Ca²⁺ signals showed arm-selective modulation, displaying sustained elevation during familiar arm traversal compared to transient novel arm responses, while astrocytes exhibited preferential activation during novel arm exploration (4.261 ± 2.172% Z-score) versus familiar arm traversal (2.663 ± 2.202% Z-score) (Fig. 6B). This differential activation was temporally organized, with astrocytic onsets occurring 600.0 ± 27.22 ms earlier than neuronal signals (Fig. S3A and S3B), suggesting their potential role in priming neuronal circuits for novelty processing.

Ischemic injury (BCAL) abolished these differential responses, reducing global Ca²⁺ activity (neurons: -0.04507 ± 0.9524% in novel arm, 0.3094 ± 0.8045% in familiar arm; astrocytes: 1.299 ± 1.139% in novel arm vs sham) (Fig. 6C, right, Fig. S3B - S3Dii) and eliminating arm-choice selectivity (novel vs familiar p > 0.05). Remarkably, hM3D-mediated astrocytic activation restored novel arm-specific neuronal Ca²⁺ signals to sham levels (novel arm: 2.210 ± 0.9669% Z-score vs BCAL -0.4100 ± 1.280% Z-score, p = 0.0058) (Fig. 6Diii and 6Div, and Fig. S3Ei), while hM4D manipulation failed to rescue these deficits (Fig. 6Diii). To further confirm that these effects were specifically driven by chemogenetic activation rather than non-specific effects of viral injection, we assessed neuronal Ca²⁺ dynamics in saline-injected control groups expressing either Gq- or Gi-coupled receptors. As shown in Supplemental Figure 1D–E, saline administration did not alter neuronal Ca²⁺ signals during novel or familiar arm choices, ruling out potential off-target effects of the chemogenetic manipulation itself.

Additionally, hM3D-mediated astrocytic activation did not enhance old arm-specific neuronal Ca²⁺ signals to sham levels (Fig. S3E). The selective rescue of novel-arm responses further supports astrocytes’ functional specialization in processing environmental novelty, potentially through modulating synaptic plasticity thresholds at information-acquisition synapses. Intriguingly, during start arm entries - regardless of whether mice transitioned from novel or familiar arms - both astrocytic and neuronal Ca^2+^ signals displayed conserved activation patterns across all experimental groups. Astrocytic signals during transitions showed no differences between sham and BCAL mice (Fig. 6D). Similarly, neuronal transition signals were comparable among sham, BCAL, hM3D, and hM4D groups (Fig. 6C). This preservation of transition-phase Ca^2+^ dynamics suggests that hippocampal network reset mechanisms may operate independently of both astrocytic modulation and prior spatial experience.

#### Gq-mediated cognitive recovery is independent of neurogenesis

Given the well-established role of hippocampal neurogenesis in cognitive function and recovery after brain injury, we next asked whether the protective effects of Gq-mediated astrocytic activation might involve alterations in neurogenesis. To address this question, we examined neural stem/progenitor cell populations, proliferation, and newborn neurons following chemogenetic activation. Immunofluorescence analyses revealed that Gq-mediated astrocytic activation did not significantly alter Sox2^+^ or Ki67^+^ cell densities in the CA1 or dentate gyrus, nor did it affect DCX^+^ newborn neuron density in the CA1 (Supplemental Fig. 4 and 5). These findings indicate that the protective effects of Gq-mediated astrocytic activation—including the restoration of cognitive function, synaptic remodeling, and precision control of hippocampal Ca^2+^ oscillations—are independent of enhanced neurogenesis.

## Discussion

PSCI represents a significant unmet clinical need, with over 30% of stroke survivors suffering persistent memory dysfunction despite modern therapeutic interventions^11, 12^. Current neuromodulation approaches, including repetitive trans-cranial magnetic stimulation (rTMS) and trans-cranial direct current stimulation (tDCS), attempt to address this challenge by broadly enhancing neuronal excitability^1, 13^. However, these strategies show variable efficacy and carry risks of seizure induction and disrupted memory consolidation due to their non-specific action on neural circuits^3, 4^. Here, our findings demonstrate that targeted chemogenetic activation of hippocampal CA1 astrocytes through the Gq pathway rescues spatial memory deficits in PSCI mice via a multi-level restorative mechanism. At the synaptic level, this intervention restored dendritic complexity, spine density, and long-term potentiation (Fig. 3 and Fig. 4). More remarkably, in vivo fiber photometry recordings revealed selective enhancement of neuronal activity specifically during novel arm exploration in the Y-maze (Fig. 6D), while maintaining baseline activity in familiar contexts. This behaviorally specific potentiation of neural coding - achieved without inducing global neuronal hyperactivation - represents a fundamental advance over conventional neuromodulation approaches like rTMS/tDCS that non-specifically enhance excitability across neural circuits. Together, these results establish a new therapeutic framework where precise astrocyte-mediated circuit modulation, rather than direct neuronal stimulation, can drive functional memory recovery after stroke.

Although astrocytes are recognized for their supportive roles in stroke—including phenotypic reprogramming (for BBB repair and neuroinflammation mitigation)^14^, lactate-dependent mitochondrial transfer to neurons^15^, NF-κB inhibition^16^, and immunomodulation via mechanisms such as BST2-C3/C3aR signaling to recruit microglia^17^—their direct regulation of post-stroke cognition remains mechanistically unresolved. Our investigation was initially inspired by the seminal work of Adamsky et al.^3^ who first demonstrated that astrocytic Gq activation enhances memory through de novo neuronal potentiation in healthy mice. We sought to extend these findings to pathological conditions by examining whether similar astrocyte-mediated mechanisms could rescue cognitive deficits after stroke. Our hypothesis gained further support from the recent work of Cho et al. ^5^ who showed that optogenetic astrocyte calcium modulation improves motor recovery in chronic stroke models. Their findings not only validated the therapeutic potential of astrocyte-targeted interventions in ischemic conditions but also encouraged our focus on cognitive-specific recovery mechanisms.

Our results reveal an important dissociation between synaptic plasticity restoration and neuronal activation patterns following astrocytic modulation. While Gq activation (hM3D + CNO) successfully rescued dendritic morphology (Fig. 3C) and long-term potentiation in CA1 neurons (Fig. 4F), it did not produce corresponding increases in c-Fos expression (Fig. 5E and F), suggesting that Gq-mediated astrocytic modulation operates primarily through synaptic-level plasticity rather than by directly increasing neuronal firing rates. This finding aligns with emerging concepts of behavioral timescale plasticity initially proposed by Bittner et al.^18^, where synaptic modifications can occur independently of immediate changes in neuronal firing rates. The preserved structural and functional plasticity without elevated c-Fos levels implies that Gq-activated astrocytes may preferentially influence the slower, sustained components of circuit reorganization that underlie memory consolidation rather than acute neuronal activation. In contrast, manipulation of the Gi pathway yielded markedly different outcomes. Despite producing comparable astrocytic activation, hM4D treatment resulted in pathological neuronal hyperactivation that exceeded sham control levels (Gi:15.42 ± 3.853% vs sham: 8.552 ± 2.955% c-Fos^+^ cells) while failing to improve cognitive performance. This dissociation likely reflects disruption of critical homeostatic functions normally maintained by astrocytes, including glutamate uptake/clearance (EAAT2), potassium buffering, and GABAergic tonic inhibition^19–21^. The resulting network destabilization appears to generate non-specific, disorganized activity patterns that are insufficient to support coherent spatial representations, as evidenced by the combination of elevated c-Fos expression with persistent behavioral impairment. These contrasting outcomes between Gq and Gi pathway modulation highlight the importance of target specificity when considering astrocyte-based interventions for neurological recovery. This conclusion is strongly supported by the work of Chai et al.^22^, who not only demonstrated fundamental differences in how hippocampal astrocytes respond to Gq versus Gi activation, with Gi signaling showing particularly weak efficacy in modulating calcium responses in hippocampus but also observed that hM4D-mediated Gi activation increased c-Fos expression in astrocytes without producing corresponding functional effects. This parallel suggests that Gi pathway engagement in astrocytes may trigger molecular activation markers (like c-Fos) while failing to induce functionally relevant circuit modulation, a phenomenon we similarly observed in our stroke model where Gi manipulation caused neuronal hyperactivation without behavioral improvement. Our findings extend these observations by demonstrating that such pathway-specific effects have direct clinical consequences for post-stroke recovery, where Gq activation promotes functional restoration while Gi engagement leads to network destabilization despite apparent neuronal activation.

Notably, this refinement exhibits behavioral state specificity. While astrocytes selectively gate novelty related neuronal Ca^2+^signals during exploration, their influence is suspended during start arm transition—a phase where both astrocytic and neuronal Ca^2+^ patterns remain invariant across all groups (Fig. 6C). This suggests hippocampal networks may alternate between two operational modes. firstly, an astrocyte-assisted state during information acquisition, where astrocytic Ca^2+^ precedes and primes neuronal plasticity. Secondly, an intrinsic reset state during trajectory initiation, where conserved dynamics override prior experience. Such dual mode operation could allow efficient memory encoding while preventing interference during navigation phase shifts.

The present study elucidates the critical role of astrocytes in modulating hippocampal Ca^2+^ dynamics during novel information encoding, particularly in the context of post-stroke recovery. We identified a remarkable temporal dissociation in CA1 activity, where astrocytic Ca^2+^ signals consistently preceded neuronal responses by 600 ms during novel arm exploration in the Y-maze. This early activation was coupled with a striking preference for novel environments, suggesting astrocytes may initiate preparatory synaptic modifications before neuronal engagement. The precise timing coincides with known windows for behavioral timescale plasticity mentioned before, implying astrocytes could lower the threshold for synaptic potentiation through targeted gliotransmitter release or metabolic support. The functional importance of this glial-neuronal coordination became particularly evident following ischemic injury. BCAL not only attenuated global Ca2+ activity but completely abolished the arm-selective responses in both cell types (Fig. 6B and 6C). A finding that mirrors the results reported by Zhou et al.^8^, who observed that astrocytic microdomain Ca²⁺ transients were significantly reduced following ischemic injury. This reduction in astrocytic Ca²⁺ activity was associated with behavioral deficits in ischemic stroke mice, highlighting the critical role of astrocytes in modulating neuronal plasticity during recovery. The selective restoration of novel arm-specific neuronal responses through hM3D-mediated astrocytic activation, while leaving familiar arm responses unaffected, which provides compelling evidence that astrocytes preferentially gate plasticity in novelty-processing pathways. The contrasting failure of hM4D-mediated Gi manipulation to improve either novel or familiar arm responses (despite inducing widespread neuronal c-Fos activation) provides further evidence that functional recovery requires precise astrocytic signaling patterns rather than general neuronal activation. Several limitations warrant consideration in interpreting our findings. While our study provides promising insights into the therapeutic potential of astrocytic Ca^2+^ signaling in stroke models, we must acknowledge the functional diversity of astrocytes - both between neural circuits and across brain regions - which underlies their multifaceted regulation of neuronal signaling and plasticity during behaviorally relevant processes^23^. More fundamentally, our chemogenetic approach targeted all CA1 astrocytes uniformly, whereas emerging evidence reveals functional micro-specialization even within this defined region. Williamson’s seminal work precisely demonstrates this principle, showing that discrete subpopulations of hippocampal astrocytes LAAS are selectively recruited during memory processes while neighboring astrocytes remain inactive^7^. This crucial finding suggests our bulk modulation approach may have simultaneously engaged multiple functionally distinct astrocyte populations, potentially obscuring more specialized roles in post-stroke recovery that could be revealed through sub population-specific interventions.

In conclusion, we demonstrate that selective Gq-pathway activation in hippocampal CA1 astrocytes rescues post-ischemic memory deficits by restoring synaptic plasticity and normalizing novelty-related neuronal Ca²⁺ dynamics. Critically, the beneficial effects are pathway-specific, as Gi activation exacerbates neuronal hyperactivation and fails to improve cognition. Our findings thus establish astrocytic Gq signaling in the hippocampal CA1 as a precise therapeutic target for cognitive recovery after stroke, and highlight the fundamental importance of pathway specificity in achieving therapeutic astrocyte modulation.

## Materials and Methods

### Animals

104 Male C57BL/6J mice (7-8 weeks, 20-23g; Guangdong Medical Laboratory Animal Centre) were group-housed under 12h light/dark cycles with ad libitum access to food/water at Zunyi Medical University. Mice were randomly assigned to groups using Excel’s RAND function. Sample size was determined via HKU Med’s calculator (α=0.05, 90% power) based on pilot stroke behavior data, indicating n = 4-5/group. Accounting for model success rate and mortality, we used n = 6-8/group. All animal procedures were performed in accordance with institutional guidelines and approved by the University Committee on Animal Research (License No. ZMU21-2302-240), following the ARRIVE guidelines for reporting in vivo experiments. Experiments were performed in a blind manner.

### Stereotactic virus injection

Mice were anesthetized with isoflurane (2.5% induction, 1-1.5% maintenance) and secured in a stereotaxic frame (RWD, China). After craniotomy, unilateral (photometry) or bilateral (chemogenetics) injections were made in the right hemisphere or both hemispheres (AP -1.94mm, ML -1.25mm, DV -1.25mm) using glass microelectrodes at 40nl/min (injection pump-controlled).

### Viral vectors

For fiber photometry, 150 nl/site of rAAV-GfaABC1D-cyto-GCaMP6f-SV40 or rAAV-hSyn-GCaMP6f-WPRE-hGH (Brain VTA, China) were injected undiluted. For chemogenetics, 150 nl/site of rAAV-GfaABC1D-mCherry-P2A-hM4D, -hM3D, or -mCherry (Brain Case, China) were similarly injected undiluted. The viruses had titers from 3.22 × 1012 to 5.59 × 1012 vg/mL.

### Transient global cerebral ischemic model

The transient global cerebral ischemia procedure was performed as previously described in our published work^10^. In brief, mice were anesthetized using isoflurane, and a midline neck incision was performed to expose both common carotid arteries. The arteries were gently encircled with surgical sutures. To induce transient cerebral ischemia, the sutures were tightened around the common carotid arteries, blocking blood flow for 60 minutes. After the ischemic period, the sutures were loosened to restore blood flow (reperfusion). Mice that did not show a reduction in cerebral blood flow of at least 75% compared to baseline levels were excluded from subsequent analysis.

### Y-maze testing combined with fibre photometry

Three days post-BCAL, mice were implanted with a fiber optic cannula (1.25 mm OD, 200 μm core, 0.39 NA) (RWD, China) positioned 0.1 mm above the viral injection site. Testing occurred in a Y-maze (30×6×15 cm) with distinct visual cues (triangle, circle, square) on each goal arm. During the first 15-min session, one arm was blocked (novel arm). After a 1-h interval, mice were reintroduced to the maze (now with all arms open) for a 5-min test session while recording astrocytic and neuronal Ca²⁺ signals. Videos were analyzed using Panlab Smart 3.0 to calculate novel arm preference (time and distance in novel arm vs. total time/distance in both goal arms).

### Clozapine N-oxide (CNO) administration

In the chemical genetics experiment, CNO (Brain Case, China) was dissolved in DMSO and then diluted in 0.9% saline to a final concentration of 0.5%. 3mg/kg CNO was intraperitoneally injected 45 minutes before the Y-maze testing.

### Immunofluorescence

Mice were transcardially perfused with ice-cold saline followed by 4% PFA. Brains were post-fixed overnight in 4% PFA (4 °C), cryoprotected in 30% sucrose (2-3 days, 4 °C), and sectioned coronally at 40 μm using a cryostat (Leica, Germany). Free-floating sections were blocked (3% NGS + 0.3% Triton X-100 in PBS, 1 h) and incubated with primary antibodies (anti-NeuN. Abcam, Brition, 1:400; anti-GFAP, Abcam, Brition 1:300; anti-c-Fos, synaptic systems, Germany,1:1000) overnight at 4 °C. After PBS washes, sections were incubated with fluorescent secondary antibodies (1:500) (Bioss, China) for 2 h at RT, followed by DAPI counterstaining.

### Golgi-Cox staining and Morphological analysis

Brains were processed using the FD Rapid GolgiStain™ Kit (FD NeuroTechnologies, INC, America). Neurons were reconstructed in Fiji’s Simple Neurite Tracer, followed by sholl analysis (8 μm concentric rings from soma to dendrite ends). Dendritic density of CA1 neurons was quantified using Fiji’s multi-point tool.

### Brain Slice preparation

Coronal hippocampal slices (400 μm) were obtained from 10–11-week-old C57BL/6J mice. After isoflurane anesthesia, brains were quickly removed, mounted, and sliced in ice-cold oxygenated sucrose-ACSF (195 mM sucrose, 2 KCl, 1.3 NaH₂PO₄, 0.2 CaCl₂, 26 NaHCO₃, 26 D-glucose) using a Vibratome. Slices were incubated in oxygenated normal ACSF (124 mM NaCl, 3 KCl, 1.25 NaH₂PO₄, 2 CaCl₂, 1 MgSO₄, 26 NaHCO₃, 10 D-glucose) at 33.8 °C for 30 min, then held at room temperature for 30 min.

### Electrophysiology

Recordings were conducted in an oxygenated normal ACSF-perfused chamber at RT. Patch electrodes (2-6 MΩ) were pulled from borosilicate glass (Sutter, America) and filled with ACSF. The Schaffer collateral pathway was stimulated using a concentric electrode (0.1-0.3mA, 0.1ms pulses at 0.05Hz) via an Iso-flex stimulator (A.M.P.I, Israel), while recordings were made from CA1 stratum radiatum using a Multiclamp 700B amplifier. Stimulation was controlled by pClamp11.2 (Molecular Devices). Test stimuli were set to evoke fEPSPs at 50% maximal amplitude. LTP was induced with 4×100Hz trains (100 pulses, 1s duration, 10s intervals). For chemogenetic experiments, high-frequency stimulation was delivered 4 min after CNO application. All recordings were performed in current-clamp mode, with data acquired through a Digidata 1550B and analyzed using Clampex 11.2 (Molecular Devices, America).

### Statistical Analysis

All statistical analyses were performed using GraphPad Prism software (version 9.0). Data normality was verified by the Shapiro-Wilk test prior to analysis. A 2-tailed t test was employed for comparison between 2 groups. For multiple comparisons, one-way ANOVA with Tukey’s post hoc test was utilized. two-way ANOVA followed by the Šídák’s post hoc test was used to compare means when there were 2 variables. Statistical significance was determined at P < 0.05. Quantitative data are reported as mean ± SD (text) or mean ± SEM (figures) throughout the study.

## Acknowledgements

This work was supported by the following funding sources:

National Natural Science Foundation of China (No.81860230)

Major Instrument Project of National Natural Science Foundation of China (No.62027812)

Guizhou Provincial Basic Research Program (Natural Science No. ZK [2024]-257)

STI2030-Major Projects 2021ZD0202900

a Ph.D. Startup Fund from Zunyi Medical university (F-878)

## Conflict of interest

The authors declare that there is no conflict of interest.

## Notes

### Competing Interest Statement

The authors have declared no competing interest.

### Summary of Updates

In this revised version, we have added a new set of results demonstrating that Gq-mediated astrocytic activation confers cognitive and synaptic protection independently of hippocampal neurogenesis. Specifically, new immunofluorescence data show that chemogenetic activation of Gq signaling in astrocytes does not alter Sox2-positive neural stem/progenitor cell densities, Ki67-positive proliferation rates, or DCX-positive newborn neuron densities in the CA1 or dentate gyrus. These findings, presented in new Supplemental Figures 4 and 5, strengthen our conclusion that the observed restoration of cognitive function, synaptic remodeling, and precision control of hippocampal calcium oscillations occur without changes in neurogenesis. No other changes were made to the manuscript.

