## supplemental materials for "Gq-pathway activation in hippocampal CA1 astrocytes rescues ischemia-induced memory deficits and synaptic plasticity"

Yujie Chen *et al.*

**This PDF file includes:**

Figs. S1 to S5

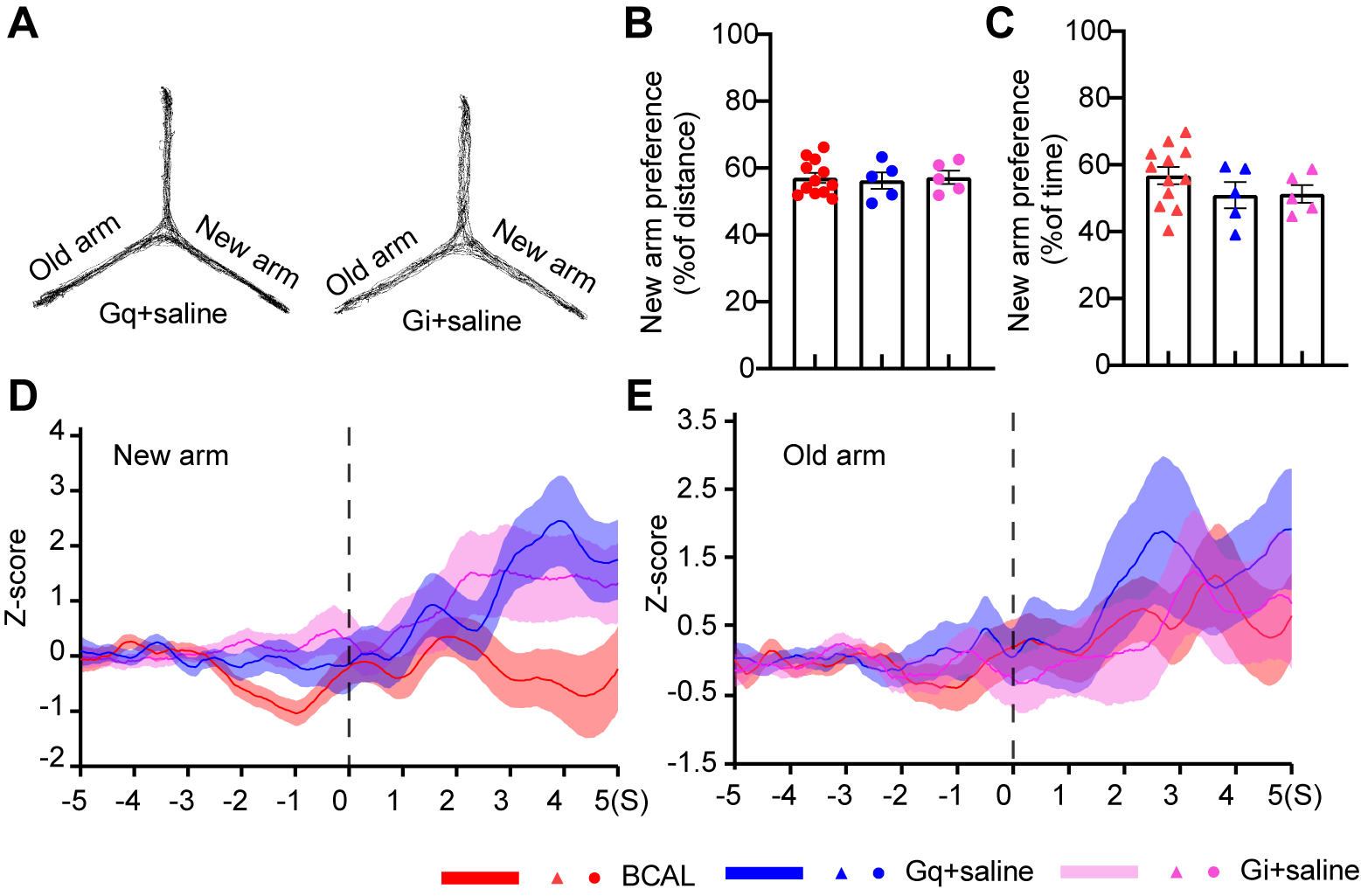

**Fig. S1.**

**Saline failing to rescue novel arm preference deficits or neuronal Ca²⁺ signaling in** **chemogenetic groups.** (**A**) Representative Y-maze traces. (**B-C**) Novel arm preference (distance (**B**) /time (**C**)) in BCAL vs Gq+saline vs Gi+saline. (**D-E**) Quantification of neuronal Ca²⁺ changes during novel (**D**) and familiar (**E**) arm choices in BCAL, Gq + saline, and Gi + saline groups. Sample sizes: n = 5 hSyn-GCaMP6f mice. Data: mean ± SEM. Statistics: one-way ANOVA.

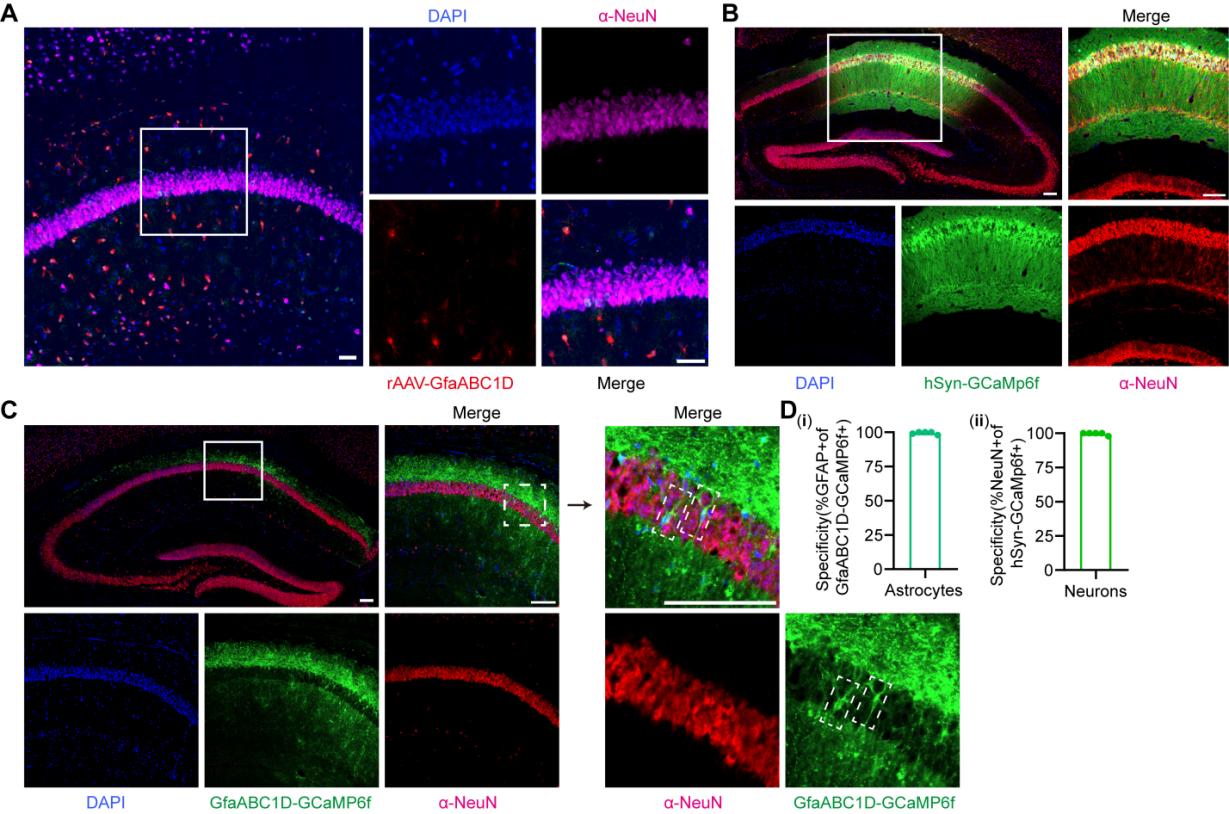

Fig. S2.

**Specific viral transduction in CA1 astrocytes and neurons.** (A-C) Confocal images demonstrating selective expression of hM4D (A), hSyn-GCaMP6f (B), and GfaABC1D-GCaMP6f (C, left) in respective cell types, colocalized with NeuN. (C, right) High-magnification view showing GfaABC1D-GCaMP6f^+^ astrocytes (white box) adjacent to CA1 neurons. (D) Quantification revealed > 97% specificity for both astrocytic (i) (410/413 cells) and neuronal (ii) (435/437 cells) expression. Data represent mean ± SEM from 5 slices/mouse. Scale bar: 100 μm.Create a page break and paste in the Figure above the caption.

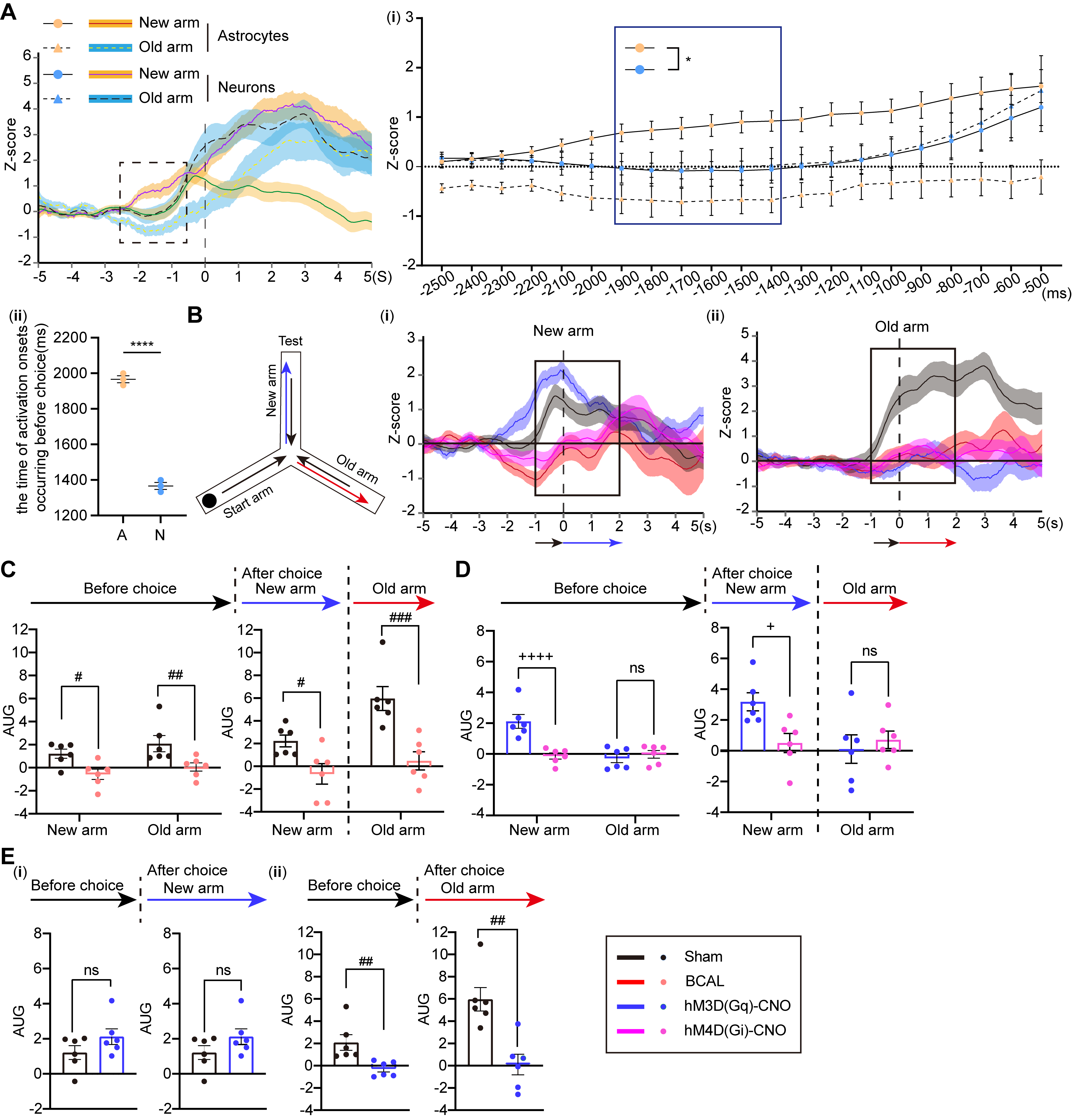

Fig. S3.

**Gq-mediated astrocytic activation potentiates neuronal Ca^2+^ responses during novel arm exploration.** (**A**) Representative Ca²⁺ transients in neurons and astrocytes during novel vs familiar arm choices in sham mice (black dotted box indicates response onset). (i) Quantitative comparison of Ca²⁺ changes between cell types (blue box highlights significant differences). (ii) Activation onset timing for astrocytes (A) and neurons (N). (**B**) Left: Y-maze schematic showing movement trajectories (black: center; red: old arm; green: new arm). Right: Typical Ca²⁺ traces during new (i) and old (ii) arm choices. (**C-D**) AUC quantification of Ca²⁺ signals in new and old arms in sham vs BCAL (C), hM3D vs hM4D (D). (**E**) Chemogenetic modulation effects in hM3D vs sham groups during new (i) and old (ii) arm exploration. Sample sizes: 9 GfaABC1D-GCaMP6f mice, 6 hSyn-GCaMP6f mice. Data: mean ± SEM. *****P* < 0.0001 vs astrocytes in new arm; ^#^*P* < 0.05, ^##^*P* < 0.01, ^###^*P* < 0.001 vs sham; ^+^*P* < 0.05, ^++++^*P* < 0.0001 vs hM3D. Statistics: two-way ANOVA. Create a page break and paste in the Figure above the caption.

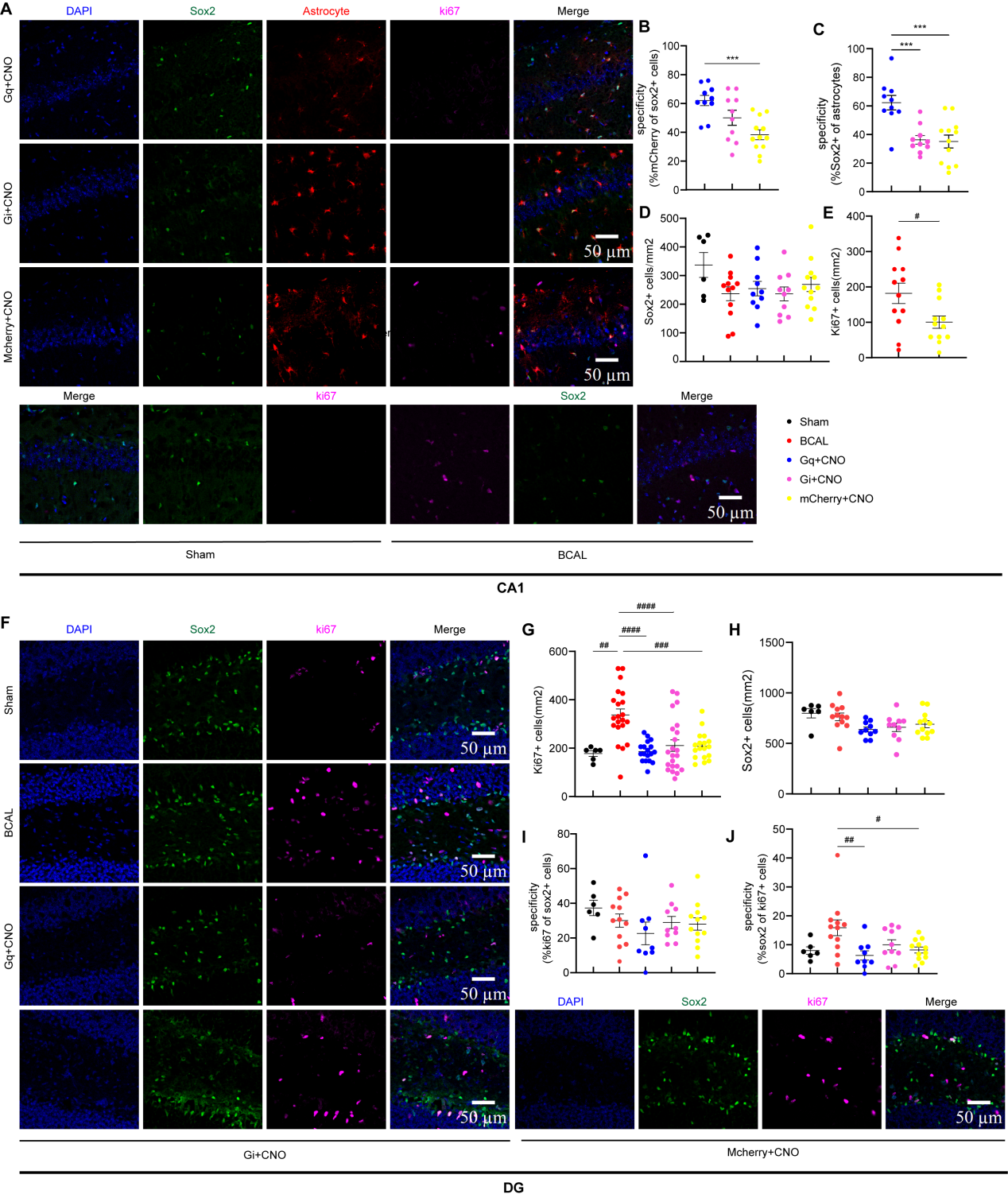

**Fig. S4.**

**Gq-mediated astrocytic activation does not alter stem cell numbers or cell proliferation.** (**A**) Representative CA1 images showing Sox2^+^/Ki67^+^ cells in sham, BCAL, chemogenetic groups. (**B-C**) chemogenetic virus (red) and Sox2 (green) in CA1 sections showing their colocalization. (**D**) Quantification of CA1 Sox2^+^ cell density in the sham, BCAL, and chemogenetic groups. (**E**). Quantification of CA1 ki67^+^ cell density in the BCAL vs mCherry group. (**F**) Representative DG images showing Sox2^+^/Ki67^+^ cells in sham vs BCAL vs chemogenetic groups. (**G**) Quantification of DG Sox2^+^ cell density in the sham, BCAL, and chemogenetic groups. (**H**) Quantification of DG ki67^+^ cell density in the sham, BCAL, and chemogenetic groups. (**I-J**) Sox2 (green) and ki67 (far-red) in DG sections showing their colocalization. Sample sizes: sham/BCAL (6-12 slices from 3-4 mice); chemogenetic groups (10-12 slices from 3-4 mice). Data: mean ± SEM. ****P* < 0.001 vs sham; ^#^*P* < 0.05 ^##^*P* < 0.01, ^###^*P* < 0.001, ^####^*P* < 0.0001 vs BCAL. Statistics: unpaired t-test (**E**); one-way ANOVA (**B**-**D** and **G-J**). Scale bar: 50 μm.

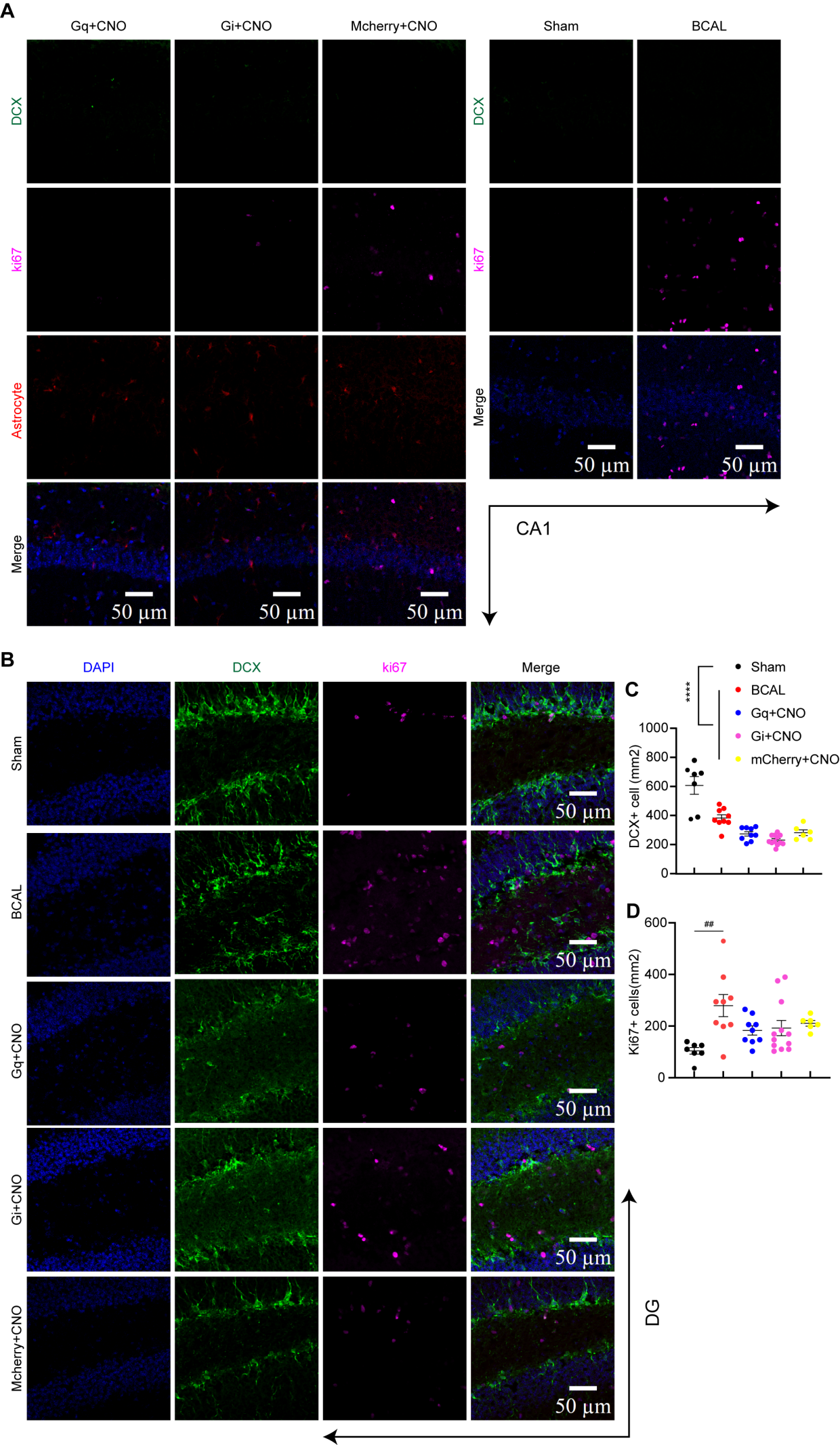

**Fig. S5.**

**Gq-mediated astrocytic activation does not promote neurogenesis.**

(**A**) Representative CA1 images showing DCX^+^/Ki67^+^ cells in sham, BCAL and chemogenetic groups. (**B**) Representative DG images showing DCX^+^/Ki67^+^ cells in sham, BCAL and chemogenetic groups. (**C**) Quantification of DG DCX^+^ cell density in the sham, BCAL, and chemogenetic groups. (**D**) Quantification of DG Ki67^+^ cell density in the sham, BCAL, and chemogenetic groups. Sample sizes: sham/BCAL (7-9 slices from 2-3 mice); chemogenetic groups (6-12 slices from 2-4 mice). Data: mean ± SEM.*****P* < 0.0001, ^##^*P* < 0.01 vs BCAL. Statistics: one-way ANOVA (**C** and **D**). Scale bar: 50 μm.
